# Optipyzer: A fast and flexible multi-species codon optimization server

**DOI:** 10.1101/2023.05.22.541759

**Authors:** Nathan LeRoy, Caleigh Roleck

**Affiliations:** Center for Public Health Genomics, University of Virginia, Charlottesville, USA; Department of Biomedical Engineering, University of Virginia, Charlottesville, USA; Systems, Synthetic, and Physical Biology PhD Program, Rice University, Houston, TX, USA

**Keywords:** codon optimization, bioinformatics, web server, recombinant protein

## Abstract

Codon optimization is a commonly used tool in many fields of scientific research to optimize the expression profile of recombinant proteins in a target organism. It seeks to swap synonymous codons in a recombinant gene to reflect the specific codon usage bias of the expression system. While many tools for codon optimization exist, they cannot optimize a single gene for expression in multiple species and are typically limited in their ability to process large sets of genes. Additionally, these tools often only provide a single interface for running optimizations, making them incompatible with existing bioinformatics tools and pipelines. To address these shortcomings we created Optipyzer: an online codon optimization web server with an accompanying web interface and Python package for multi-species codon optimizations. Optipyzer was designed to be fast, flexible, and extensible - providing numerous interfaces for query submission. We believe that Optipyzer can be a powerful tool for the modern biologist by seamlessly integrating into any workflow. The main web interface can be accessed at https://optipyzer.com.

## Introduction

Recombinant protein expression is an important biotechnological tool used in various fields including medicine, agriculture, food production, manufacturing, and scientific research. (1)(2)(3)(4). Across all areas, a common challenge is to optimize the expression profile of these recombinant proteins in a new target organism. One approach is to optimize mRNA folding energies post-translation, which has been shown to be effective (5). However, this is not the only method. One common cause of low expression levels of engineered recombinant proteins is a reduction in the translational efficiency due to variances in codon preference between the origin and target organisms (6). For this reason, many researchers employ codon optimization to swap synonymous codons in a recombinant gene to reflect the specific codon usage bias of the expression system. Codon optimization increases translational efficiency by ensuring that the recombinant protein uses codons with cognate tRNAs that are highly available, thus increasing the overall protein expression (7). While codon optimization might not be suitable for all applications, and while the exact mechanism and efficacy of codon optimizations are still not fully understood, the practice is still widely used today (3).

While many computational tools for codon optimization exist (8) (9) (10) (11) (12) (13), there are two limitations with current approaches: 1.) No existing algorithm or tool can optimize a sequence for expression in multiple species, despite its potential benefits for applications including synthetic ecology (14) (15), *in situ* microbiome engineering (16), and comparative studies on performance of a single genetic construct across multiple species (17); 2.) present tools especially those which are web-based - are limited in their ability to process large sets of genes. To this end, we set out to create a multispecies codon optimization algorithm that can not only optimize a given gene for multiple species, but one that can do so in a high throughput manner with speed, security, and flexibility in mind. Here we present Optipyzer: A multi-species codon optimization server with an accompanying web interface and Python package for flexible optimization workflows. Optipyzer leverages the most up-to-date codon usage data through the HIVE-Codon Usage Tables database to ensure an accurate representation of the specific codon usage bias of any species. (18) (Fig. 1A).

**Fig. 1.**
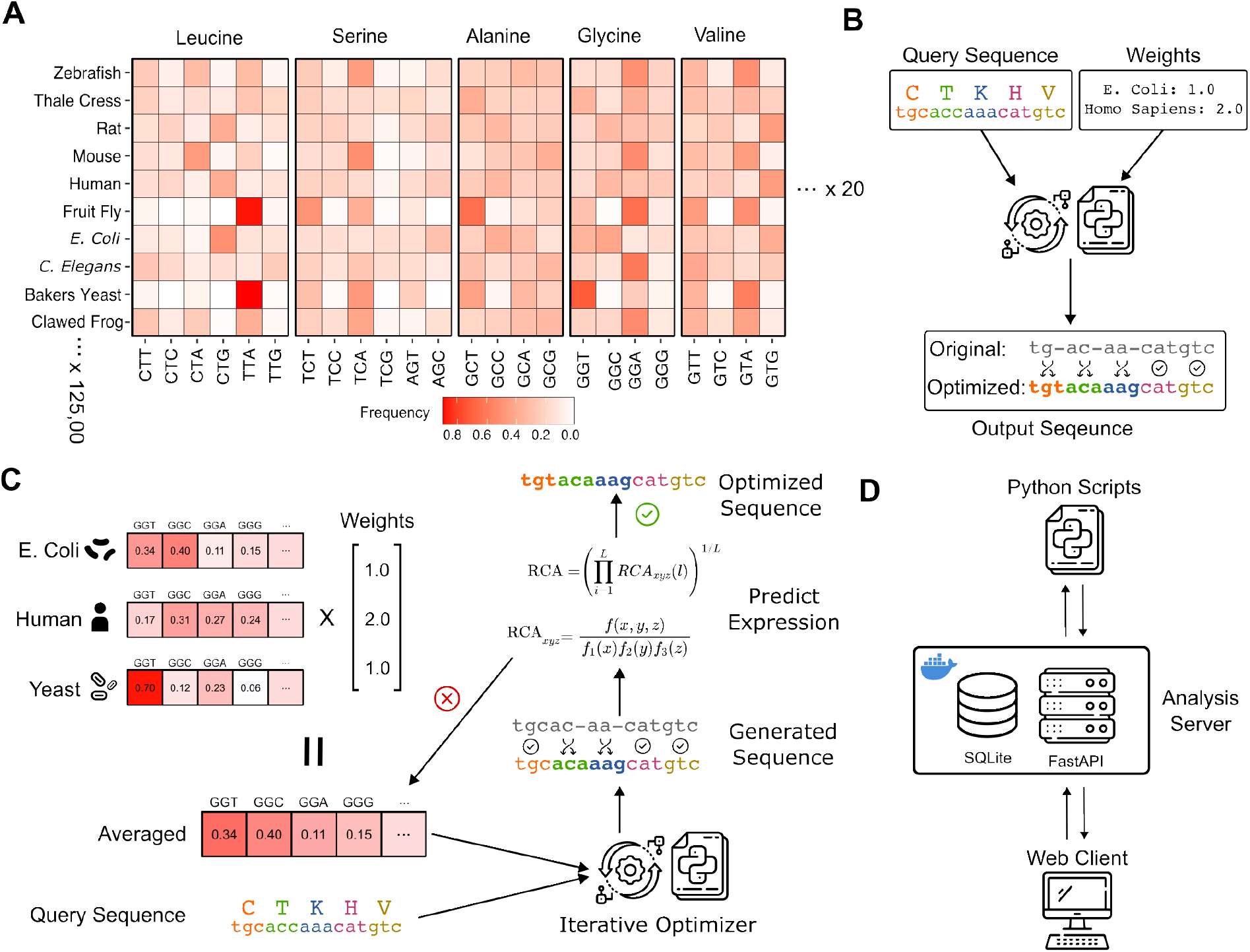
**a.)** Heatmap describing the codon usage preferences for the top 24 most common codons and 10 popular species. Each species uses codons in a preferential manner. **b.)** Basic input and output flow of Optipyzer. A query sequence and species weights are required as input, and Optipyzer will output an optimized sequence. **c.)** Detailed flow of the Optipyzer algorithm. Species codon usage data are combined using a weighted average to create a unified usage table. This table is combined with a query sequence and a new sequence is generated. The new sequence is checked using RCA to predict the relative expression in each species. If the relative expression does not align with the input query, the combined table is updated and a new sequence is generated and checked again. Once the sequence exhibits the desired predicted expression profile the algorithm stops. **d.)** The architecture of Optipyzer is structured with a central optimization server. The web client and Python API communicate to the server through HTTP.

## Methods

### The Algorithm

Optipyzer consists of a three-step algorithm. First, the codon usage data for each organism in the query is pulled from the database. Second, an averaged table is computed using each species’ individual tables and the species weights from the query via a weighted average. Finally, the averaged table is used to construct an optimized query using a stochastic selection process and the relative codon adaptation index (RCA) to ensure a proper expression profile (Fig. 1C).

#### Step 1

In step one we pull a codon usage table for each species in the query from the HIVE-CUTs (19) database. These tables are based on the frequency that codons appear in protein-coding regions of the genome and contain raw counts. The counts are then pre-normalized to a codon usage bias value between 0 and 1.

#### Step 2

Step two consists of the average table calculation. The averaged table is constructed by calculating a weighted average for each codon frequency using the species weights provided in the query:

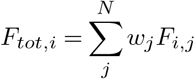

*F*_*tot,i*_ is the total weighted frequency for codon *i, w*_*j*_ is the species weighting in the query and *F*_*ij*_ is the specific species usage frequency for codon *i* in species *j*.

This process might result in the calculation of rare codons. A rare codon is defined as any codon that codes for its respective amino acid less than 10% of the time (20)(21). Codons considered rare for any species in the optimization are thus considered “prohibited” with a frequency of 0 for all species in the optimization, and the non-prohibited codons are renormalized so that the sum of the frequencies per residue adds to one.

#### Step 3

Finally in step three, an optimized sequence is generated using the “codon randomization” method of optimization (22) by sampling codons according to the frequencies defined in the averaged table. To confirm that our resulting sequence is optimized to the specified weighting during optimization, we employ the use of a relative codon adaptation index (RCA), which has been shown to correlate to protein abundance data,(23) to confirm that the measure of relative expression for each species is consistent with the original query. The RCA index is calculated with the following formula:

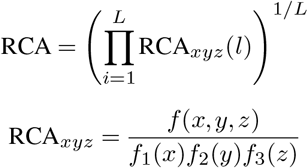

*f* (*x, y, z*) is the observed frequency of codon *xyz* and *f*_1_(*x*), *f*_2_(*y*) and *f*_3_(*z*) the observed frequencies of bases *x, y* and *z* at, respectively, codon positions 1, 2 and 3 in the reference genome and *L* is the number of codons in the gene to be optimized. (24).

To determine whether or not the predicted expression matches the desired expression ratio across the input species, the RCA values are normalized so that the lowest RCA value is set to 1. Both the absolute value and the sum of the squares of the difference between the target and predicted expression values are calculated. If the optimized sequence does not exhibit a predicted expression ratio that matches the desired input in the query, the averaged frequency table is updated by slightly adjusting the weights of six codons in favor of species which currently have an expression difference greater than 5% of its target expression value. A new optimized sequence is generated and the sum of squares and absolute difference of predicted and target expression values are calculated and compared to that of the previously generated sequences to determine the best performing sequence. This process is iterated upon 1000 times to ensure that the optimized sequence exhibits the best expression profile possible for all species in the query in regards to codon bias and irrespective of other factors that might contribute to expression differences across species.

## Results

### Open-source web API

*Optipyzer* was designed to function as a microservice that can be integrated with any workflow and improve interoperability of molecular biological pipelines (25) (Fig. 1D). At its core, *Optipyzer* is a RESTful web API that accepts optimization queries and computes the result on a remote server. Any application or client can utilize HTTP to interact with *Optipyzer*, allowing it to fit many workflows. To our knowledge, this is the first codon optimization platform that implements a web API. Furthermore, *Optipyzer* is open-sourced to embody the power of collaboration and scientific reproducibility (26). The majority of current codon optimization tools - especially those of which are web-based - are not open-source. This hinders transparency, communication, and collaboration. *Optipyzer* enables these core tenets through its open-source and distributed nature (Fig. 2). API documentation can be found at https://optipyzer-api.herokuapp.com/docs

**Fig. 2.**
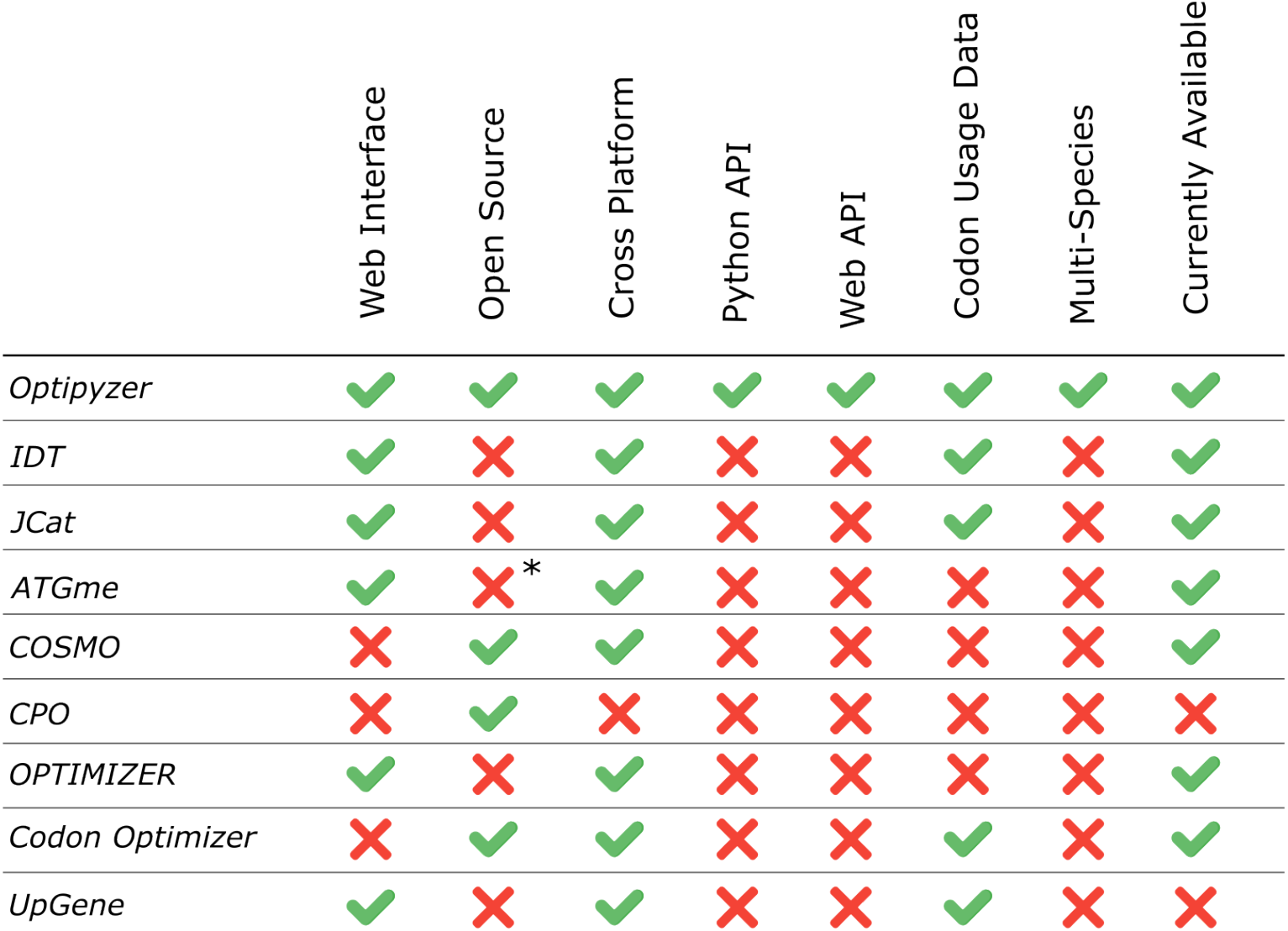
Feature comparison. Typically, codon optimization tools are not open-sourced (IDT, JCat, ATGme). If they are (COSMO, CPO, Codon Optimizer), they lack an architecture that facilitates web-based interaction through graphical or programmatic means. Further, no current method implements a web API which is critical for integration with other tools and platforms. \**ATGme* is reported as open-source (21), but no links to source code could be found.

### Python Package

All optimization requests are made through the web service. While this facilitates flexibility, we wanted to make it as easy as possible for researchers to utilize the web API. To this end, we also developed an accompanying Python package that provides a convenient API to utilize *Optipyzer* from within Python programs and scripts. Users can install the package from PyPi using `pip` and begin utilizing *Optipyzer* programmatically. Further, since requests are made through HTTP, any language that maintains an HTTP library can also leverage the service. (E.g. *R*: `httr`, *Perl*: `HTTP::Request`, *JavaScript*: `fetch()`). Because we maintain our own codon usage database (18) on the server, to our knowledge, *Optipyzer* is also the first python-based codon optimization package that doesn’t require the tedious and manual curation of codon usage tables. View the python package on GitHub: https://github.com/nleroy917/optipyzer

### Web Interface

We also have implemented a web-based graphical user interface. Accessible via web browser, users can construct, submit, and view optimization requests using a convenient web application. The web interface also leverages our web API, further demonstrating the ability of *Optipyzer* to be flexible and be used for many diverse applications. During this process, users can also view codon usage statistics for any organism. Users input their sequence, select organisms from a drop-down, and specify optimization hyperparameters (random number generator seeds and number of iterations). To our knowledge, *Optipyzer* is also the first open-source codon optimization web application.^1^ The web application can be found at https://optipyzer.com.

### Example Use

*Optipyzer* requires two inputs: 1.) The DNA or protein query in FASTA format; and 2.) a list of organisms and their respective targeted relative expression levels (Fig. 1B). An example query is given below in JSON:

~~~
{
   “seq”:”tgcaccaaacatgt…”,
   “weights”: {
      “human”: “1”,
      “e_coli”: “2”
   }
}
~~~

This example query is optimizing a nucleotide sequence for expression in both *Escherichia coli* and *Homo Sapiens* cell lines with a 2:1 weighting towards human cell lines. We can then use `curl` to submit the optimization request. This simple demo exemplifies *Optipyzer’s* ease of use and potential for integration into any existing tool.

~~~
curl -XPOST -H “Content-type: application/json” -d \
‘{
      “seq”: “tgcaccaaacatgttgcaccaaacatg”,
      “weights”: {
        “e_coli”: “1”,
        “human”: “2”
     }
}’ \
‘https://optipyzer-api.herokuapp.com/optimize/dna‘
~~~

### Additional Web Resources

The Optipyzer architecture is modular (Fig. 1D). It facilitates many different client types, is extensible, and scales. The core of Optipyzer is a web server implemented in Python using the FastAPI framework. Since the codon usage database is sufficiently small (45 Mb) and infrequently updated, it is packaged entirely alongside the web server as a SQLite database instance. The web server has endpoints capable of querying the database for the codon usage data of any species, searching for species that exist in the database, and also running optimizations. For detailed information on endpoint availability and usage, visit https://optipyzer-api.herokuapp.com/docs.

The web-based client is implemented in React.js using the Next.js framework. The client leverages the optimization endpoints to optimize sequences from queries that users make on the site. The site makes query submission as convenient as possible by providing text boxes, species search fields, and dropdowns to construct queries. Users can also view codon usage statistics for any individual organism. Optimization requests are made through HTTP.

For more advanced or high-throughput workflows, a Python interface was created and is pip installable through PyPi. This way, users can write their own scripts natively in Python to optimize sequences that may be a part of other pipelines or in files held locally on their machine. In the interest of both speed and security, we also prepared a Docker container to run an instance of the optimization server with codon usage data included locally. This drastically improves optimization time and improves security as all requests never leave the host machine.

## Conclusions

Optipyzer is a new fast and effective multi-species codon optimization server capable of optimizing recombinant DNA sequences for multiple target organisms simultaneously. It includes a web server for running optimization queries, a web-based client for running queries, and a Python API to integrate codon optimization into custom scripts and workflows. While not yet experimentally validated, we believe that Optipyzer will give researchers a flexible tool to best generate, profile, and characterize any protein they want in as many organisms they want.

## ACKNOWLEDGEMENTS

We gratefully acknowledge the editorial support of Dr. Nathan Sheffield and project guidance from Dr. Michael Gribskov.

## Footnotes

1 *ATGme* is reported as open-source (21), but no links to source code could be found.

